# Memory-driven computing accelerates genomic data processing

**DOI:** 10.1101/519579

**Authors:** Matthias Becker, Milind Chabbi, Stefanie Warnat-Herresthal, Kathrin Klee, Jonas Schulte-Schrepping, Pawel Biernat, Patrick Günther, Kevin Baβler, Rocky Craig, Hartmut Schultze, Sharad Singhal, Thomas Ulas, Joachim L. Schultze

**Affiliations:** Platform for Single Cell Genomics and Epigenomics, German Center for Neurodegenerative Diseases (DZNE) and the University of Bonn, Bonn, Germany; Genomics and Immunoregulation, Life & Medical Sciences (LIMES) Institute, University of Bonn, Bonn, Germany; Hewlett Packard Labs, Palo Alto, USA; Hewlett Packard Enterprise, Ratingen, Germany

**Keywords:** memory-driven computing, genomics, RNA-sequencing, ecosystem, systems biology

## Abstract

Next generation sequencing (NGS) is the driving force behind precision medicine and is revolutionizing most, if not all, areas of the life sciences. Particularly when targeting the major common diseases, an exponential growth of NGS data is foreseen for the next decades. This enormous increase of NGS data and the need to process the data quickly for real-world applications requires to rethink our current compute infrastructures. Here we provide evidence that memory-driven computing (MDC), a novel memory-centric hardware architecture, is an attractive alternative to current processor-centric compute infrastructures. To illustrate how MDC can change NGS data handling, we used RNA-seq assembly and pseudoalignment followed by quantification as two first examples. Adapting transcriptome assembly pipelines for MDC reduced compute time by 5.9-fold for the first step (SAMtools). Even more impressive, pseudoalignment by near-optimal probabilistic RNA-seq quantification (kallisto) was accelerated by more than two orders of magnitude with identical accuracy and indicated 66% reduced energy consumption. One billion RNA-seq reads were processed in just 92 seconds. Clearly, MDC simultaneously reduces data processing time and energy consumption. Together with the MDC-inherent solutions for local data privacy, a new compute model can be projected pushing large scale NGS data processing and primary data analytics closer to the edge by directly combining high-end sequencers with local MDC, thereby also reducing movement of large raw data to central cloud storage. We further envision that other data-rich areas will similarly benefit from this new memory-centric compute architecture.

---

Next generation sequencing (NGS) and its manifold applications are transforming the life and medical sciences^1,2,3,4^ as reflected by an exponential 860-fold increase in published data sets during the last 10 years (Fig. 1a). Considering only human genome sequencing, the increase in genomic data over the next decade has been estimated to reach more than three Exabyte (3×10^18^ byte)^5^. NGS data are projected to become on par either with data collections such as in astronomy or YouTube or the most demanding in terms of data acquisition, storage, distribution, and analysis^5^. In addition, since one of the cornerstones of genomics research and application is comparative analysis between different samples, long-term storage of genomic data will be mandatory. To cope with this data avalanche, so far, compute power is simply increased by building larger high-performance computing (HPC) clusters (more processors/cores), and by scaling out and centralizing data storage (cloud solutions)^6,7^. While currently still legitimate, it can already be foreseen that this approach is not sustainable. Centralized storage and computing leads to increased data movement, data duplication, and additional risks to data security and privacy. This is of particular interest given recent data privacy laws that have been put in place (e.g. the General Data Protection Regulation (GDPR) in the European Union)^8^. Another important aspect that requires re-thinking algorithms and the required compute infrastructure to cope with the growing requirements of genomic research is the enormous energy consumption of data centers providing the respective cloud services. Moreover, since many current algorithms are not scaling well with increasing data volumes, just increasing (scaling out) classical HPC infrastructures might not suffice. A solution to these emerging limitations requires reduction of 1) compute time, 2) data movement, 3) data duplications, and 4) energy foot print to a minimum, while increasing 5) data and privacy protection at the same time. We addressed these issues by combining a novel computer hardware architecture – namely memory-driven computing (MDC) – with optimized algorithms in genomic data analytics.

**Fig. 1:**
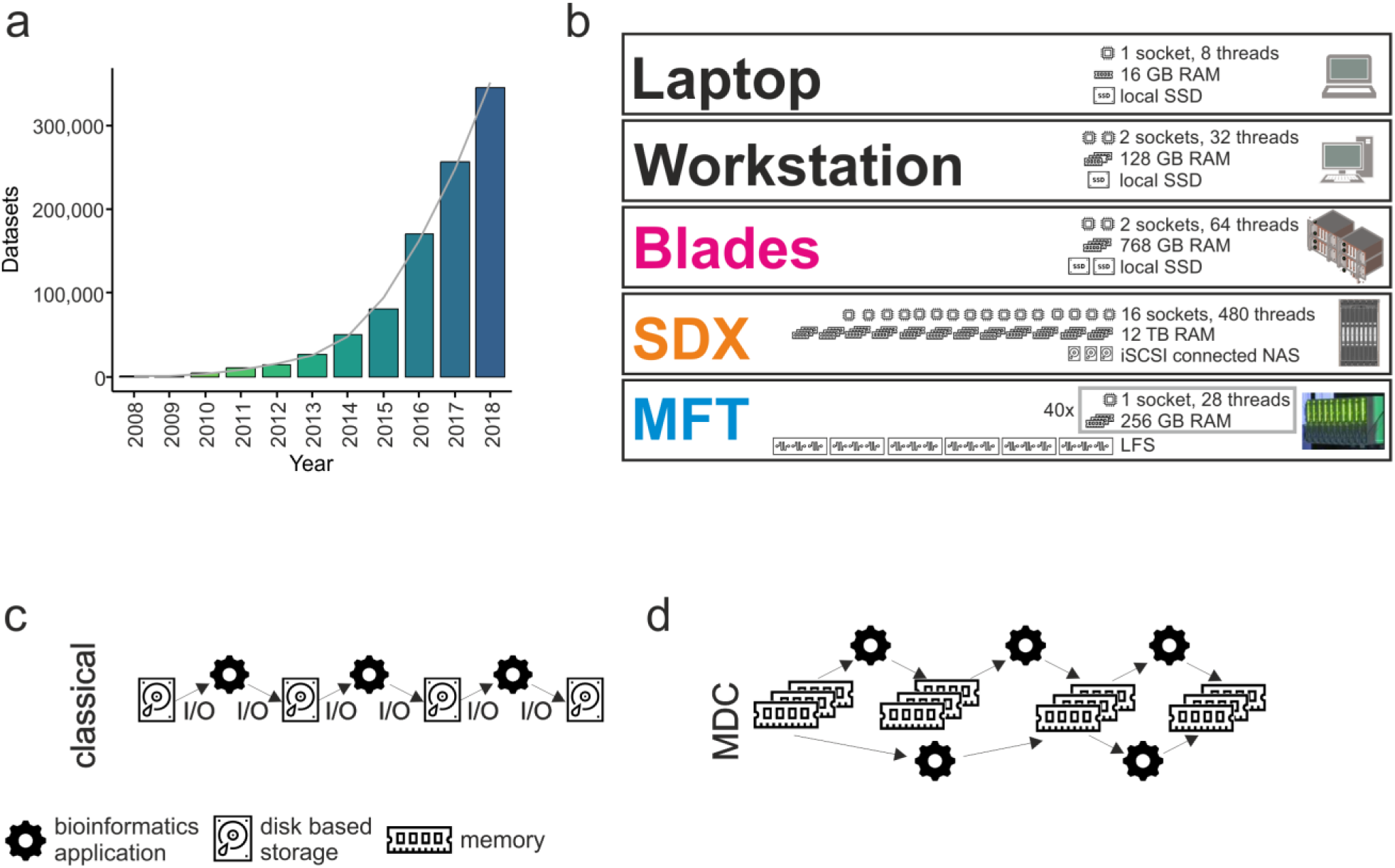
Adaptation of genomic data processing to memory-driven computing. **a,** Depiction of genomic datasets published since 2008. **b,** Different hardware infrastructures and their technical specifications as they have been used here. **c, d,** Schematic representation of data handling within classical computing infrastructures with sequential I/O handling for each task (**c**) and within MDC allowing processing of data completely available in abundant faster memory (**d**). RAM, random access memory; SSD, solid state drive; iSCSI, internet small computer system interface; NAS, network attached storage; LFS, librarian file system; I/O,

All classical computers, from laptops to super computers, are based on the von Neumann architecture from 1945^9^. As of 2018, a typical single user system (Laptop/Desktop) has 1 CPU with 4 to 8 CPU-Cores with typically 8 to 16GB of RAM. Workstations typically have 2 CPUs with 64 to 128GB of RAM (Fig. 1b) although up to 3 TB are possible. Multipurpose single server systems start in similar ranges and scale higher in CPU-Cores (up to 224) and memory (up to 12TB). Servers (standalone, rackmount or blade systems) may be clustered over high-performance interconnects to scale CPU-resources and workloads may be deployed on the cluster using a scheduler. Specialized HPC systems may reduce space requirements further by higher integration. Nowadays, scale up interconnected systems can provide up to 48TB of main memory acting as a single server instance. In contrast to the von Neumann architecture (Extended Data Fig. 1a), MDC can pool large, shared and persistent (storage class) memory and connects this memory and compute resources (CPU) via a fabric following the Gen-Z protocol (‘Fabric Attached Memory’, FAM) (Extended Data Fig. 1b)^10^. While traditional systems are processor-centric (Extended Data Fig. 1a), in MDC the FAM is the central component (Extended Data Fig. 1b) thereby avoiding I/O (Extended Data Fig. 1c) by allowing all processors to directly access the FAM (Extended Data Fig. 1d). The MDC architecture enables all processors in the cluster to operate on shared data concurrently and the combination of MDC and the Gen-Z protocol provides the necessary byte-addressable scaling up to 4096 Yottabyte (2^92^ byte) of FAM (Extended Data Fig. 1e). Yet, long-term storage or archiving can still be accessed via traditional I/O-handling (Extended Data Fig. 1f). MDC has been realized as a prototype system called the Memory Fabric Testbed (MFT) with 160 TB of memory, but the concept can also be emulated on classic architecture using a FAM Emulation (FAME) software^11^. It has been hypothesized that processing time and compute power can be substantially reduced by eliminating the need for I/O handling within the application^12^. The use of so-called ‘in-memory’ processing systems has been a first attempt towards this direction, however, unlike MDC, these systems do not offer large persistent memory and still require fetching and storing data for processing from I/O-based storage systems.

Genomic data processing pipelines often consist of serial tasks. Using classical hardware infrastructure, each task loads small parts of a large data set from storage to local volatile memory (RAM) and writes its results back (Fig. 1c). Subsequent tasks need to fetch the data again, incurring high file access overhead^13^. With MDC, all data can be held in memory (Fig. 1d) and accessed as necessary throughout the pipeline, thus avoiding the penalty associated with moving data to disk between pipeline stages. Two main adaptations are needed to move applications to MDC: per-application optimization (Extended Data Fig. 2a) and pipeline-wide optimization (Extended Data Fig. 2b). First, individual applications can be modified for more efficient data access, and the abundance of memory enables in-memory data structures to be optimized for processing speed. Second, data exchange between parts of the pipeline can be optimized by not only reading from memory but also using in-memory data formats that reduce costly data transformations and parsing across the processing pipeline.

**Fig. 2:**
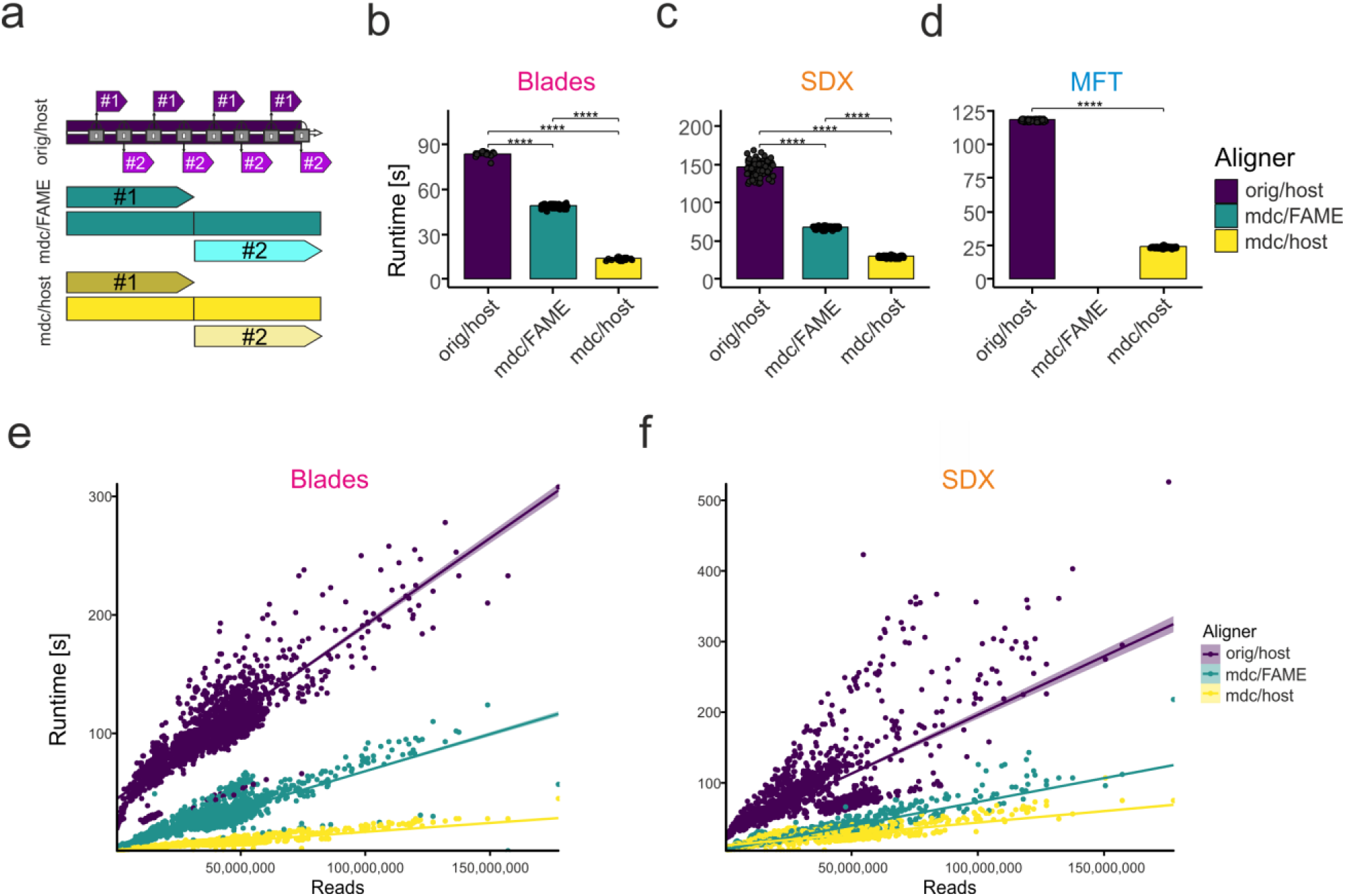
Performance analysis of RNA-seq read processing using classical and MDC-based systems. **a,** Schematic representation of the changes in data access and data size introduced to kallisto using the MDC-version either directly on the host or within an emulation environment (FAME). **b, c, d,** Runtimes of a 140 million read RNA-seq data sample using the original kallisto, or the MDC-version either directly on the host or in an emulation environment assessed on Blades (**b**), the SDX (**c**), or the memory fabric test bed (**d**; MFT). **e, f,** Assessment of runtimes of 1.181 RNA-seq samples of different read sizes on Blades (**e**) or the SDX (**f**) using the same aligner as described in 2b-c. orig/host, original kallisto code run directly on the respective hardware; mdc/FAME, MDC-adapted kallisto code run within the FAME environment on the respective hardware; mdc/host, MDC-adapted kallisto code run directly on the respective hardware.

Irrespective of source, all NGS data require lengthy pre-processing by either alignment (including pseudoalignment) to an annotated reference or de novo assembly prior RNA-seq quantification and further data analysis^14^, ^15^. To demonstrate the benefits of MDC for RNA-seq alignment and quantification, we used one of the fastest methods introduced as near-optimal probabilistic RNA-seq quantification (kallisto)^16^. Currently, kallisto reads small chunks of data in a serial fashion and processes them in parallel (Fig. 2a, orig/host). In contrast, the modified version of kallisto overcomes the data access bottleneck by allowing random parallel access and processing of larger data packages (Fig. 2a, mdc/FAME, mdc/host). The modified version of kallisto can be executed directly on classical systems with sufficiently large memory (RAM) or via the FAME emulation environment (for further technical details see Methods). We examined our modifications using the RNA-seq dataset (ERR188140) with 140 × 10^6^ reads introduced recently^16^ (Fig. 2b, c, d) on two systems for the classical architecture (Blades, Superdome X (SDX), Fig 2b, c) and the MFT for the MDC architecture (Fig. 1d, Extended Data Fig. 3). Our results clearly show that the modifications of kallisto significantly reduced the runtime within each of the tested systems (Fig. 2b, c, d). Not unexpectedly, emulation does not produce the same gain as native hardware, but even under these conditions MDC-modified code showed a significant gain over the original. To determine general applicability and scaling, we collected 1,181 publicly available RNA-seq data sets derived from blood donors with different diseases from a total of 19 studies (Extended Data Fig. 4). To rule out any influence of the underlying hardware, we used both the Blades and the SDX environment. All the RNA-seq data sets were processed with both the original kallisto version and the MDC-modified version either within the FAME environment or directly on the system (Fig. 2e, f). Independent of the size of the individual dataset, the MDC-modified algorithm always outperformed the original version both within FAME and directly on hardware resulting in reduced runtime of up to 13.4-fold. Even more surprising the gain of performance was disproportionally higher with larger data size. We further assessed kernel and user times (Methods) and found kernel time to be negligible (data not shown) while user time revealed a reduction of 66% by the MDC-modified version of kallisto when executed directly on the Blades or the SDX (Extended Data Fig. 5a, b). Also considering parallelism of data processing we observed an up to 112-fold reduction of compute time in comparison to the original hardware and software settings. To determine accuracy between the different versions of the algorithm, we compared the RNA-seq results derived by the original version of kallisto, the MDC-modified version and two alternative aligners (HiSat2, STAR) (Fig. 3a). MDC modification to kallisto only showed very minor differences (correlation 1-7.5×10^−9^), while we observed lower correlations with HiSat2 and STAR, a finding that was previously reported^16^.

**Fig. 3:**
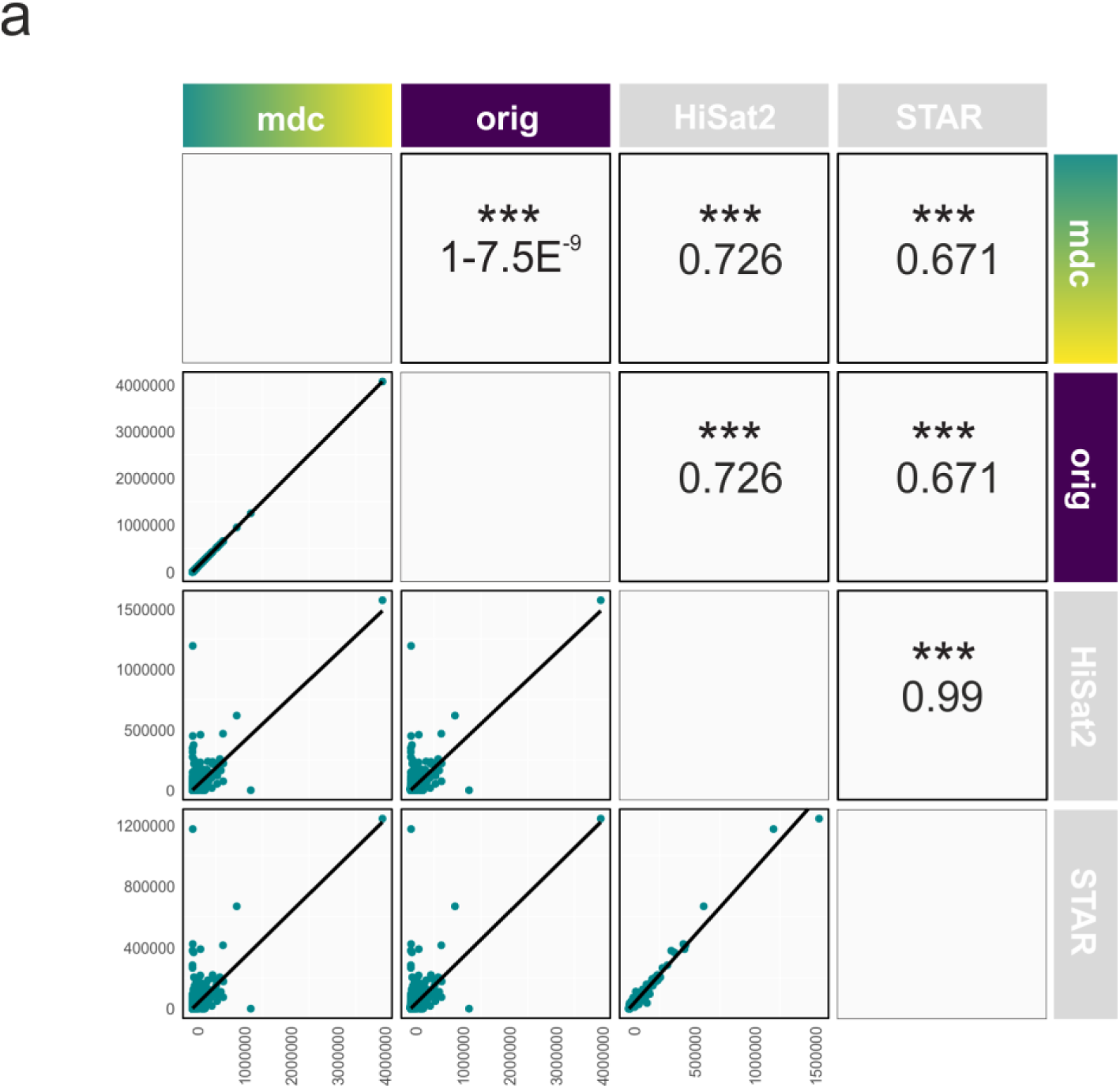
Comparison of RNA-seq quantification results from different aligners. **a,** Scatter plots comparing two aligners together with black trend lines and respective mean correlation coefficient for quantified raw gene counts from the sample ERR140188 for the algorithms MDC-modified kallisto (mdc), original kallisto (orig), HiSat2 and STAR (left to right). Pearson correlations and significance for comparisons between two aligners are shown above the diagonal. Axis depict raw gene counts.

**Fig. 4:**
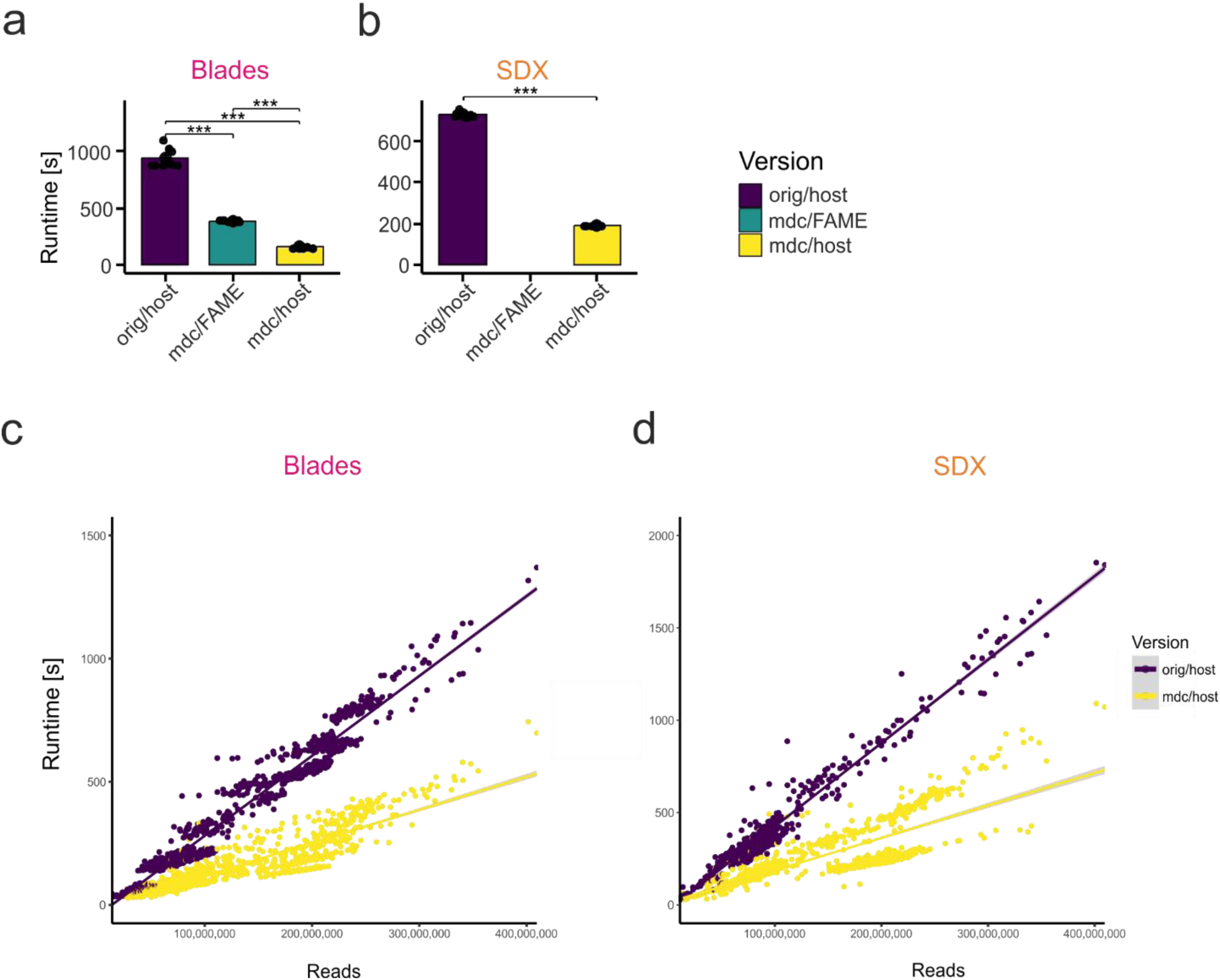
Performance comparison of NGS data assembly using a classical and a MDC-based system. **a, b,** Runtime of the RNA assembly pipeline’s first step (SAMtools) for the ERR188140 data set (**a**) and on the SDX (**b**). **c, d,** Assembly (SAMtools) runtimes for the 1.181 RNA-seq samples of different read sizes on the Blades (**c**) and the SDX (**d**). orig/host, original kallisto code run directly on the respective hardware; mdc/FAME, MDC-adapted kallisto code run within the FAME environment on the respective hardware; mdc/host, MDC-adapted kallisto code run directly on the respective hardware.

We have chosen transcriptome assembly to demonstrate the effect of pipeline wide optimizations (for detailed technical description see methods). We followed the steps proposed by Pertea et al.^17^ and have focused on SAMtools^18^ and StringTie^19^ that run serially with I/O based data exchange (Extended Data Fig. 2b). Prior work by Hajj et al.^20^ demonstrated that in-memory processing can accelerate SAMtools. We modified the sorting function to load and sort data in parallel without any temporary files and to keep the results in memory for StringTie to avoid disk-based data exchange. Our initial findings on SAMtools show a significant acceleration of up to 5.9-fold on the Blades and SDX for the ERR188140 data set (Fig. 4a, b). These finding were supported by the larger 1181 sample data set (Fig. 4c, d). Future work will focus on addressing the current bottlenecks in StringTie.

Collectively, genomic research and usage of genomic data for precision medicine require significant reduction of compute time, data movement, data duplication and energy consumption. We provide compelling evidence that these goals can be reached by moving genomic data processing to MDC, as exemplified for RNA-seq assembly and pseudoalignment. Considering the increase of memory also allowing for an increase of parallelism of data processing we demonstrate an up to 112-fold reduction in compute time in comparison to the previously used single core system for probably one of the most sophisticated RNA-seq data quantification strategies available to date (kallisto)^16^. Within one computer system, the modifications of kallisto resulted in up to 13.4-fold reduction of runtime and up to 66% reduction of user time (Methods), which is an indicator for reduced energy consumption. Similarly, a significant runtime reduction was also observed for the first step of the assembly of RNA-seq data. We predict that similar results for data processing can be obtained for all other NGS-based data (e.g. genome and epigenome alignment and assembly) even up to the single cell level. Our results clearly indicate that we can gain major benefits by investing in both optimized data analytics algorithms as well as the underlying hardware and software infrastructure. MDC seems to be an ideal architecture model since it accommodates the requirements of future computing in biomedical research. MDC is centered on data rather than processors (Extended Data Figure 1b); is modular due to the memory fabric specifications (Gen-Z) (Extended Data Fig. 6a); allows the integration of specialized analysis hardware^21^; can be connected to local producers of large data (Extended Data Fig. 6b); is scalable from Gigabytes to Petabytes and more (Extended Data Figure 1e); can reduce energy consumption (Extended Data Fig. 5a, b, c); and allows the integration of concepts of edge computing and federated cloud computing^22,23^ (Extended Data Fig. 6c). In genomics research MDC would allow handling of raw data in close proximity to sequencing devices (edge computing) (Extended Data Fig. 6b). Such MDC-supported edge computing of genomics data would still enable the development of federated cloud computing with the important difference that large genomic raw data would be kept locally at the edge where the data are produced, while inferred data could be shared within a federated (cloud) system of edge MDC devices (Extended Data Fig. 6c). Clearly, data movement and duplication would also be reduced significantly, since only smaller inferred data would be shared between different sites. Instead, algorithms could be designed to interrogate data kept at the different edges and collect inferred data from different sites in larger studies. And lastly local storage of genomic raw data at the data producers’ sites using MDC devices would reduce data privacy, security, and sovereignty issues particularly in medical applications, since such data would not have to be distributed, duplicated or stored at central cloud storage any more.

## Acknowledgements

This work was funded in part by the HGF grant sparse2big, the FASTGenomics grant of the German Federal Ministry for Economic Affairs and Energy and the EU project SYSCID under grant number 733100. We thank Bill Hayes, Keith Packard, Patrick Demichel, Binoy Arnold, Robert Peter Haddad, Eric Wu, Chris Kirby for supporting us in running the experiments on the HPE infrastructure. JLS is member of the excellence cluster ImmunoSensation.

## Methods

### Theoretical background to Memory-Driven Computing (MDC)

#### MDC and FAME in comparison to current systems

In Memory-Driven Computing (MDC), memory is the central component of the computing system (Extended Data Fig. 1b), which is in contrast to classical hardware infrastructures that are processer-centric (Extended Data Fig. 1a). In MDC, memory is accessible via a fabric and computational resources, like CPUs, GPUs or specialized processors can be attached to it. In 2018, the Gen-Z consortium has formulated standards for this fabric^10^. Unlike in classical hardware, remote memory is presented as ‘local’ to the processor and can be accessed directly through the memory controller of the processor without the overhead of layers like operating systems or network stacks (Extended Data Fig. 1c, d). Optimized memory controllers to fully embrace MDC are still in development. However, systems with large amounts of memory exist and can be used to emulate this environment. Furthermore, Hewlett Packard Labs have built a prototype, the Memory Fabric Test bed (MFT), with the largest single memory of 160 TB, which was also available for testing the performance of genomic data processing algorithms after adaptation to the MDC infrastructure. The FAME (fabric attached memory emulation) emulator^11^ provides programming abstractions that enable development of MDC tools and frameworks. It emulates an MDC environment by setting up several virtual machines that all share a large memory pool and provides direct memory access to shared memory across operating system instances running in the different virtual machines^11^.

MDC is not to be confused with so-called ‘in-memory’ approaches. Within the last two decades several such approaches (e.g. Apache Ignite Memory-Centric Storage, Hazelcast IMDG® In-Memory Data Grid, Memcached, Redis to name a few) have been developed with the same intention, namely to increase overall performance. However, all these in-memory implementations build upon standard systems based on the von Neumann compute architecture, where the CPU acts as the gateway for in-memory operations and as such is limited to the given memory bandwidth and maximum addressable memory size between the CPU and its attached memory (DRAM, NVM, SCM). Conceptually, the above-mentioned implementations circumvent the boundary of the maximum addressable memory size given by the von Neumann architecture on an application level while still being limited for speed by the maximum bandwidth given.

#### Addressing data access and internal data structures by MDC

In this work, we have assessed two key approaches in genomic data analysis, namely assembly and pseudoalignment– here exemplified for the transcriptome. Technically, we provide evidence for two typical computing approaches within such analysis that are benefiting from MDC. First, we can change the data access patterns. This is important for pipeline-wide optimizations with data exchange between tools in the pipelines through memory in custom format instead of conversion followed by writing to disk and a subsequent requirement for conversion. A second possibility to benefit from MDC is per-application optimization where we can exploit the abundance of memory to optimize constrained internal data structures. We describe both concepts in more detail in the following sections.

#### Data access changes for pipeline-wide optimizations

In classical compute infrastructures, a major bottle neck for large data is the transfer between the storage system and processing nodes. Furthermore, large data does often not fit into single nodes and needs to be partitioned. Once data is stored in abundant memory, random and parallel access to large data files is no longer subject to the massive performance penalty imposed by the necessity for I/O that is known from traditional data storage systems. Within the MDC architecture, one possibility for data access and management is controlled by a service called Librarian that divides the pool into shelves and books and is a low-level interface for allocation and access. Working with the librarian is possible through multiple ways. To allow existing applications to easily benefit from MDC, the memory pool is exposed as a virtual file system, the Librarian File System (LFS). A first step would be using just the LFS instead of traditional storage, but to fully benefit from LFS, the contents of the file need to be memory-mapped (mmap). Mmap presents file contents as a continuous region of memory and replaces typical I/O operations. This allows the operating system to efficiently load the file data asynchronously. LFS is optimized in such a way that it only needs to provide the corresponding memory pages from the complete memory pool. In the future, working with the librarian will also be possible via an API that provides just a thin abstraction layer^24^.

In both our use cases (assembly and pseudoalignment of NGS data), we have modified the index / reference loading code to use mmap. The chosen pseudoalignment and quantification algorithm (kallisto) is characterized by a data loading model that relies on a single thread that reads the data in chunks and distributes these chunks to worker threads that perform the actual matching to the reference genome. This model requires synchronization between reader and workers (Fig. 2a, top). This overhead is small for few workers but becomes a bottleneck for higher worker numbers that are possible on systems like the Blades or the SuperDomeX. We have modified kallisto and removed the separate reader process. Each worker now has its own memory region of the input assigned to (Fig. 2a, middle, bottom). This was possible by realizing the input directly in the LFS where random access carries no penalty. As the NGS assembly use case we used the combination of SAMtools and StringTie^18^, ^19^. In this case, we changed the SAMtools in a way that the input process makes use of mmap thereby allowing parallel data processing. To further accelerate the interaction between SAMtools and StringTie, we replaced the intermediate SAM file and instead used a memory-backed file for sharing. This change not only removes I/O handling but also allowed to store the SAM records in a format that can be directly leveraged by StringTie without further need to parse an input file.

#### Internal data structures optimization per application

A second method to benefit from MDC is the exploitation of the abundance of memory for internal data structures.

In the pseudoalignment use case, we recognized that kallisto uses a static hash map. This hash map allows fast access to reference genome information using a k-mer as the key. Our analysis of this process showed that lookup operations in this hash table consume most of the time in the pseudoalignment phase. Hash maps use slots to store the data. With fewer slots (higher load factor), there are collisions and additional comparisons are needed to find a specific element. With a lower load factor, the number of collisions is reduced (Extended Data Fig. 7a) at the cost of increased memory consumption. Clearly, with MDC we can make use of the abundance of memory to overcome this bottleneck. By decreasing the load factor of the hash map and hence increasing the hash map from 2 GB to 50 GB we removed most of the collisions (Extended Data Fig. 7a). Furthermore, utilizing mmap and the LFS to load the hash map – despite the fact that the index file was increased to 50 GB – was still faster than loading the hash map within the original kallisto (Extended Data Fig. 7b). Since the index file is on LFS, it can be shared between multiple instances.

For the assembly uses case, the optimization of internal data structure was achieved by removing intermediate files for data exchange. The SAMtools pipeline uses temporary files stored on slow data storage devices such as hard drives or network storage when sorting the reads in an input SAM file by either query name or genomic coordinate. With our modifications, the input is loaded concurrently in small chunks into memory. With MDC and its abundance of memory, the whole input and output data can be stored in fast (and persistent) memory. The loaded subsets are then pre-sorted and later merged. Since all these steps take place in memory no I/O handling is required anymore.

### Experimental setup addressing the benefits of MDC

#### RNA-seq data sets utilized in this study

As a first test dataset the same RNA-seq data previously described as test dataset (ERR188140) to evaluate kallisto was obtained from the Gene Expression Omnibus (GEO) database^16^. To assess performance increase in a real-world big data scenario, we compiled an RNA-seq dataset containing 1.181 transcriptomes derived from peripheral blood mononuclear cell (PBMC) samples collected within 23 different studies (GSE97309) (Extended Data Fig. 4). The majority of samples were derived from patients with Acute Myeloid Leukemia (AML; 508), followed by juvenile idiopathic arthritis (JIA; 132), vaccination against Tuberculosis (TB.vaccination; 119) as well as numerous other diseases, but also from healthy individuals (healthy; 65). All data sets in this study were published in the National Center for Biotechnology Information Gene Expression Omnibus (GEO, Barrett et al., 2012). Basic criteria for inclusion were the cell type under study (human peripheral blood mononuclear cells (PMBCs) and/or bone marrow samples) as well as the species (Homo sapiens). Furthermore, we excluded GEO SuperSeries to avoid duplicated samples. We filtered the datasets for high-throughput RNA sequencing (RNA-seq) and excluded studies with very small sample sizes (< 10 samples for RNA-seq data). We then applied a disease-specific search, in which we filtered for acute myeloid leukemia, other leukemia and healthy or non-leukemia-related samples. The results of this search strategy were then internally reviewed and data were excluded based on the following criteria: (i) exclusion of duplicated samples, (ii) exclusion of studies that sorted single cell types (e.g. T cells or B cells) prior to gene expression profiling, (iii) exclusion of studies with inaccessible data. Other than that, no studies were excluded from our analysis. All raw data files were downloaded from GEO.

#### Hardware platforms used for evaluation and performance experiments

A key goal of this work is to show that the optimization of hardware and software environments by moving towards MDC-based architectures can accelerate life science computational tasks while increasing energy efficiency. Furthermore, we also intended to demonstrate that making use of the principles of MDC on classical hardware systems already accelerates genomic data processing, irrespective of the hardware used. For our evaluations, we used three different systems: 1) a typical blade server system that is common for small computational clusters, 2) an HPE SuperDome X (SDX), a powerful system with a large number of cores and an abundance of memory that we envision to be more typical for memory centric architectures, and 3) the Memory Fabric Testbed (MFT) developed at Hewlett Packard Labs, an experimental prototype that demonstrates the principles of Memory-Driven Computing using independent processors that can share memory over a fabric.

All three systems are non-uniform memory access (NUMA) systems. NUMA describes architectures that are made from interconnected building blocks (nodes), e.g. a single CPU with many cores and its directly connected memory can form a node. These individual nodes can collaborate using a fast interconnect bus. The MFT nodes have local (exclusive to the node) memory and additionally access to a shared memory pool instead of traditional storage. This pool can be local or reachable via the memory fabric, although this difference is hidden to the programmer.

The blade system used consisted of two NUMA nodes with a total of 32 cores or 64 threads and 768 GB of DDR4 SDRAM (double data rate fourth-generation synchronous dynamic random-access memory) interface (Extended Data Fig. 3a). SSDs (Solid State Disks, 2 TB) were used for mass storage. The SDX used had 16 NUMA nodes with a total of 240 cores or 480 threads, 12 TB of DDR3 SDRAM and a storage system connected via iSCSI (Extended Data Fig. 3b). The prototype MFT used consisted of 40 nodes that share a 160 TB memory pool. One major difference between the Blades and the SDX system used in our setup is memory bandwidth. The Blades are connected with the fast DDR4 interface while the SDX is connected via the previous generation of memory interface (DDR3) only. This difference in memory bandwidth must be considered during the interpretation of the performance analysis of our highly memory dependent applications. Unlike the other systems, the MFT is not based on the x86 architecture but on ARMv8 and comes with a custom memory controller to enable MDC. Each node houses 4 TB of memory pool and 256 GB local memory (Extended Data Fig. 3c). Each CPU has 28 cores.

#### Calculation of runtime

We have used the Linux command line utility “time” to measure the runtime time for our experiments. Time is a wrapper around the program call and outputs different values, including runtime (as measured by an independent clock on the wall) as well as user and system time (see below).

#### Configuration of the pipelines

The pseudoalignment and quantification use case is realized by calling kallisto in the pseudoalignment mode (pseudo) with an estimated average fragment length of 200 with a standard deviation of 20. We used the Gencode hg38 v27 as index and single end reads as input.

The data preparation for the assembly pipeline was performed using fastp^25^ for cleanup of the input files and Hisat2 for producing SAM files. Then SAMtools were used for sorting and StringTie for reference guided assembly. Fastp was called with parameters for a minimum phred score of 15, an unqualified base limit of 40%, per read cutting by quality score in front and tail, tail trimming and poly-G/X trimming with a minimum length of 10. For the remaining tools of the assembly pipeline, we followed the protocol by Pertea et al.^17^.

#### Comparison of outputs

Reads in FASTQ format from the test sample ERR188140 were quantified at the transcript level against an Ensembl catalog using the original and the modified version of kallisto, aggregated to the gene level using tximport and the genome annotation database org.Hs.eg.db v3.6.0 in R. These 25.541 raw kallisto counts were combined with the 19.320 raw read counts coming from HiSat2 and STAR. Genes were only kept if they were identified by all three algorithms resulting in a matrix of 17.731 unique genes (Fig. 3). The correlation analysis was performed using the ggpairs function from the package GGally v1.4.0, which is an extension of ggplot2.

For the second use case, assembly, we verified the correctness of the results using gffcompare^26^. We used gffcompare with a reference annotation (Gencode hg38 v27) and compared each sample against it. After completing the runs on the different platforms, we found identical results for all conditions tested (data not shown). This result was expected since the changes to the assembly pipeline were designed to modify data access without altering the inner workings of the assembly algorithm, in particular within StringTie.

#### Impact on energy consumption

Computer operating systems are organized in layers, central and hardware related functions (e.g. I/O) are placed in the kernel space, with kernel being the core component of the operating system. Application software and their related functionality are executed in user space and can, if required, access kernel functionality, for example to fetch data (Extended Data Fig. 5d, e). If a program is parallelized, the total user time from a parallel component is the sum of user time of each thread (*t*_*user*_*=*Σ_*p*∈ *threads*_ *t*_*user*_(*p*)) and often larger than the wall clock (processing) time. In our application, the user time is the main contributor to the total runtime. The user time often grows with the number of threads. However, with good parallelization and less overhead in data access the user time might be lower as user threads might not be waiting for the kernel (e.g. from I/O operations which are computationally expensive).

In the pseudoalignment use case, we have measured the runtime (Fig. 2e, f), which can be used to calculate the total saving of time. Since these savings have been reached with a larger number of threads, we have also looked at the user and kernel times. These values include the sum of all threads and show how long each corresponding core was used in total. Kernel time was found to be negligible in the kallisto use case. The comparison of user time values revealed a reduction of 66% by the MDC-version of kallisto when executed directly on the SDX or the Blades (Extended Data Fig. 5b, c). In the FAME emulation environment, the reductions are lower, due to virtualization overheads, but the user time is still lower. The MFT results (Extended Data Fig. 5a) are similar to the overall 66% user time savings that we found in the MDC-version of kallisto. The user time can be considered an indicator for energy consumptions and points to possible savings with MDC.

#### Data security and access control in MDC

Albeit we have not directly evaluated aspects of data security and access control in our experiments, we would like to point out that MDC offers additional benefits over classical computing infrastructures concerning these important issues. Genomics data are most sensitive concerning data privacy and therefore, we need to pay special attention to securing such data. This is of particular importance for systems such as MDC where the shared memory pool is envisioned to be persistent and can enable access to that data over time to different programs and users. Not surprisingly, data access control^27^ and data security^28^ have been integral parts of the design of the Gen-Z fabric connecting all components^10^. An implicit functionality of MDC and Gen-Z is the logical isolation of resources to prevent unauthorized access to sensitive data and the support of authenticated and encrypted communication between components. Benchmarking this functionality against current solutions will require further future research.

#### Kallisto new version comparison

During the preparation of this manuscript, a new version of kallisto (v0.44) has been released. We inspected the code changes for differences in data access patterns and parallelism and found no changes for these parts of the code. Nevertheless, to verify that the changes did not impact on our results we tested the runtime of the original (v0.43.1) and the updated (0.44) version of kallisto on a test data set and found no significant changes (Extended Data Fig. 8). We concluded from this comparison that a re-analysis of our results with the newer version of kallisto was not required for answering the questions proposed in this project.

### Data availability

The data sets used in this work can be found at the Sequence Read Archive (ERR188140) and the Gene Expression Omnibus data base (1181 samples, GSE97309). The modified source code of kallisto and SAMtools can be found on GitHub at https://github.com/LIMES-immunogenomics/.

**Extended Data Fig. 1:**
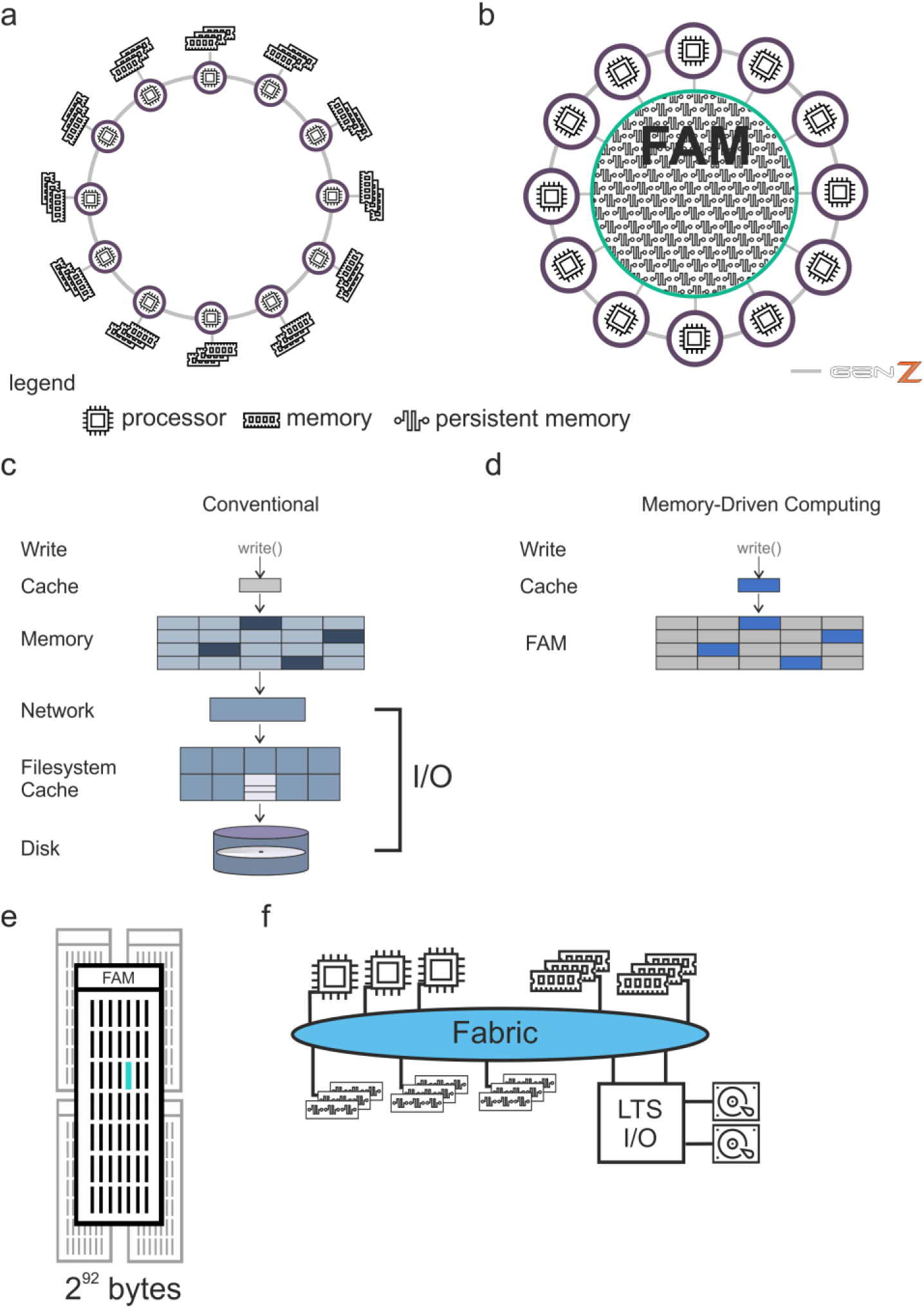
Memory-driven computing (MDC) background. **a, b,** Depiction of traditional cluster architecture (**a**) and MDC architecture (**b**). **c, d,** Layers needed to access storage in traditional system (**c**) and in the MDC architecture (**d**). **e,** The MDC architecture provides byte-addressable scaling to large persistent storage-classed memory pools that act like a single system. **f,** Long-term storage (LTS) and archive can be accessed via traditional I/O-handling.

**Extended Data Fig. 2:**
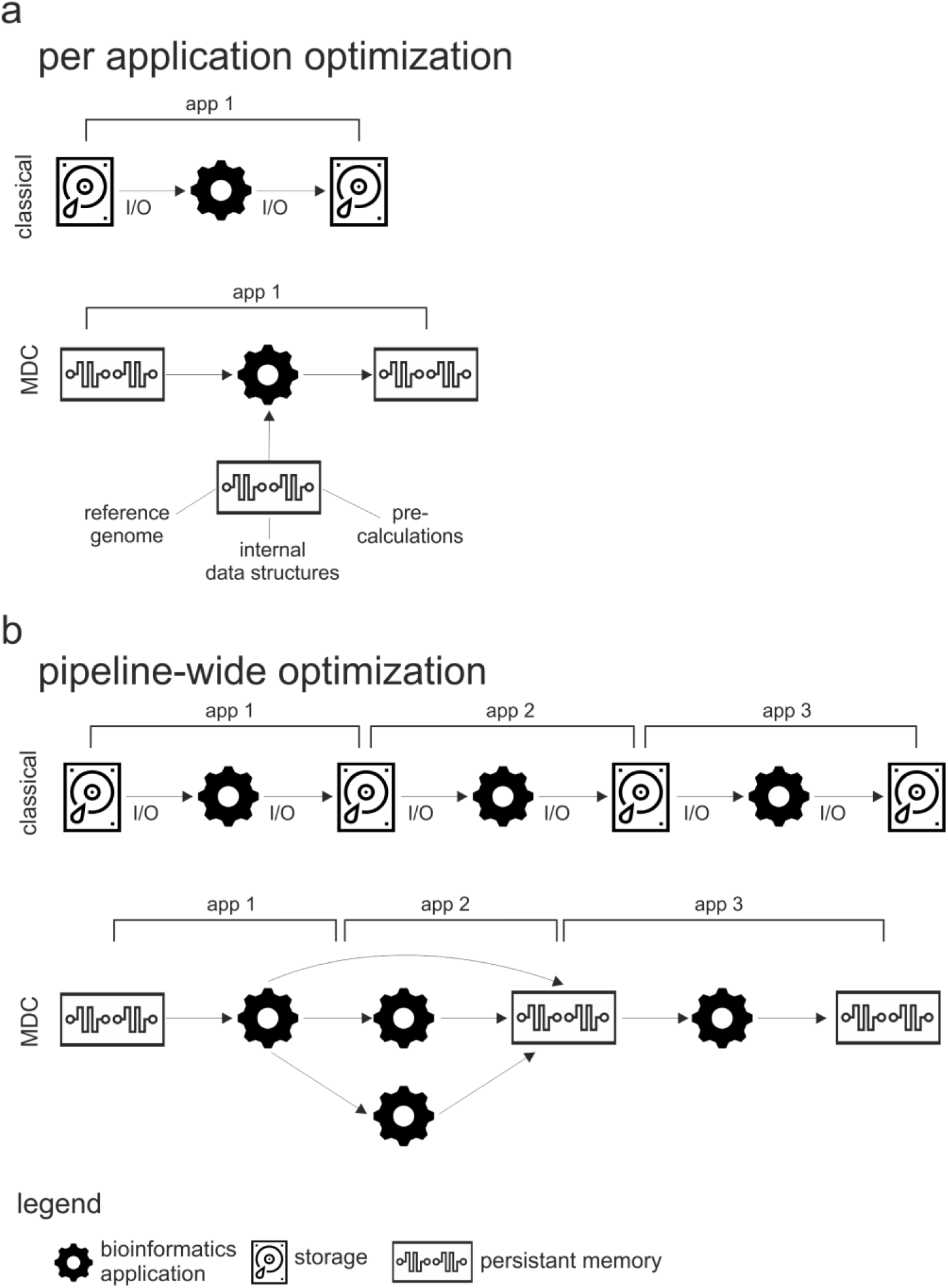
Optimization approaches. **a,** Classical single-tool pipeline with I/O handling and access to disc storage systems and MDC optimization by accessing all data directly from memory. **b,** Classical multi-tool pipeline with read and write operations via I/O on disc storage systems and MDC enabled pipeline-wide optimizations.

**Extended Data Fig. 3:**
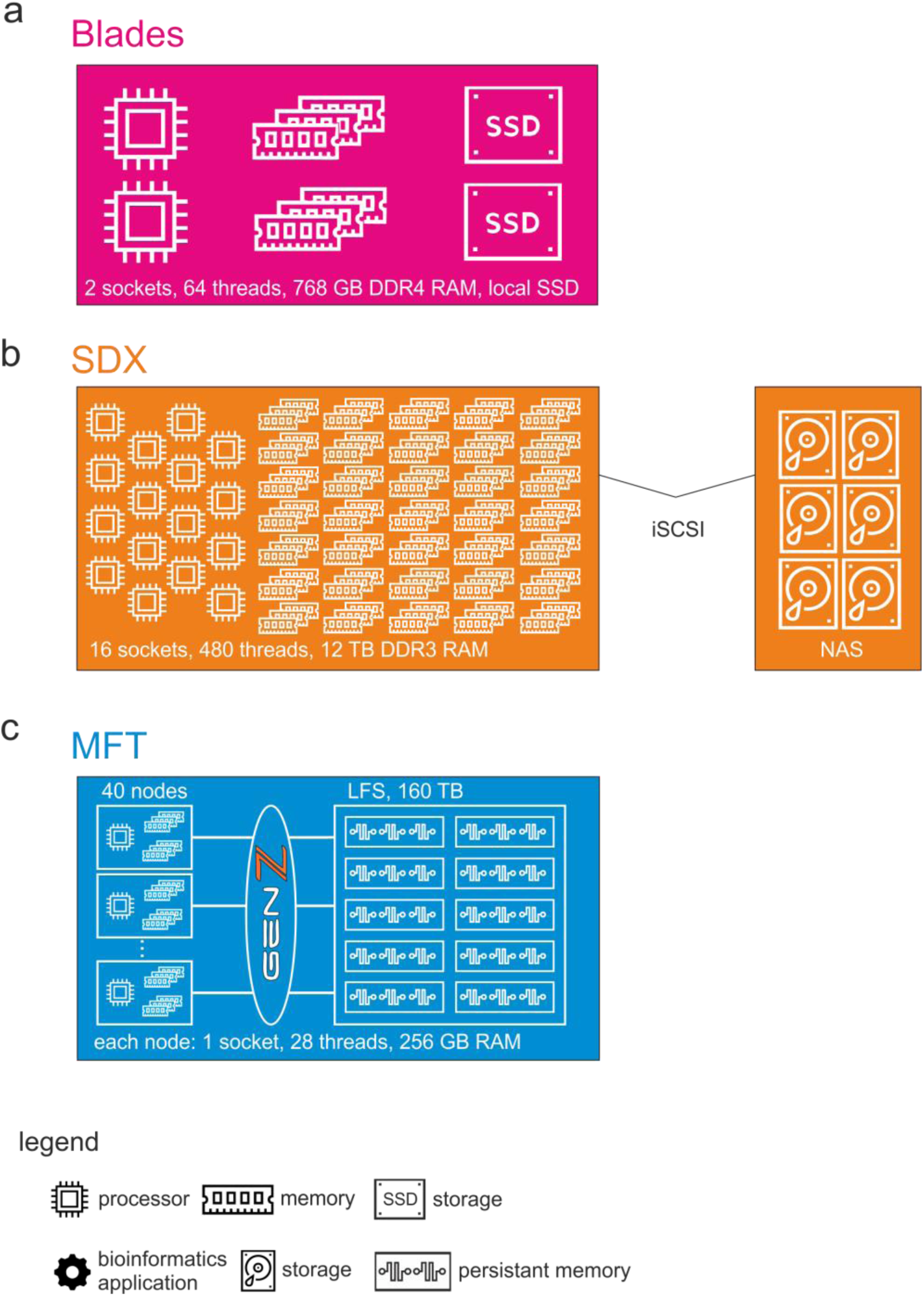
Systems used for evaluation. **a,** Blade system with two sockets and local SSD storage. **b,** Large server with fifteen sockets and remote disk-based storage and **c,** the memory fabric testbed with two sockets and access to the memory pool.

**Extended Data Fig. 4:**
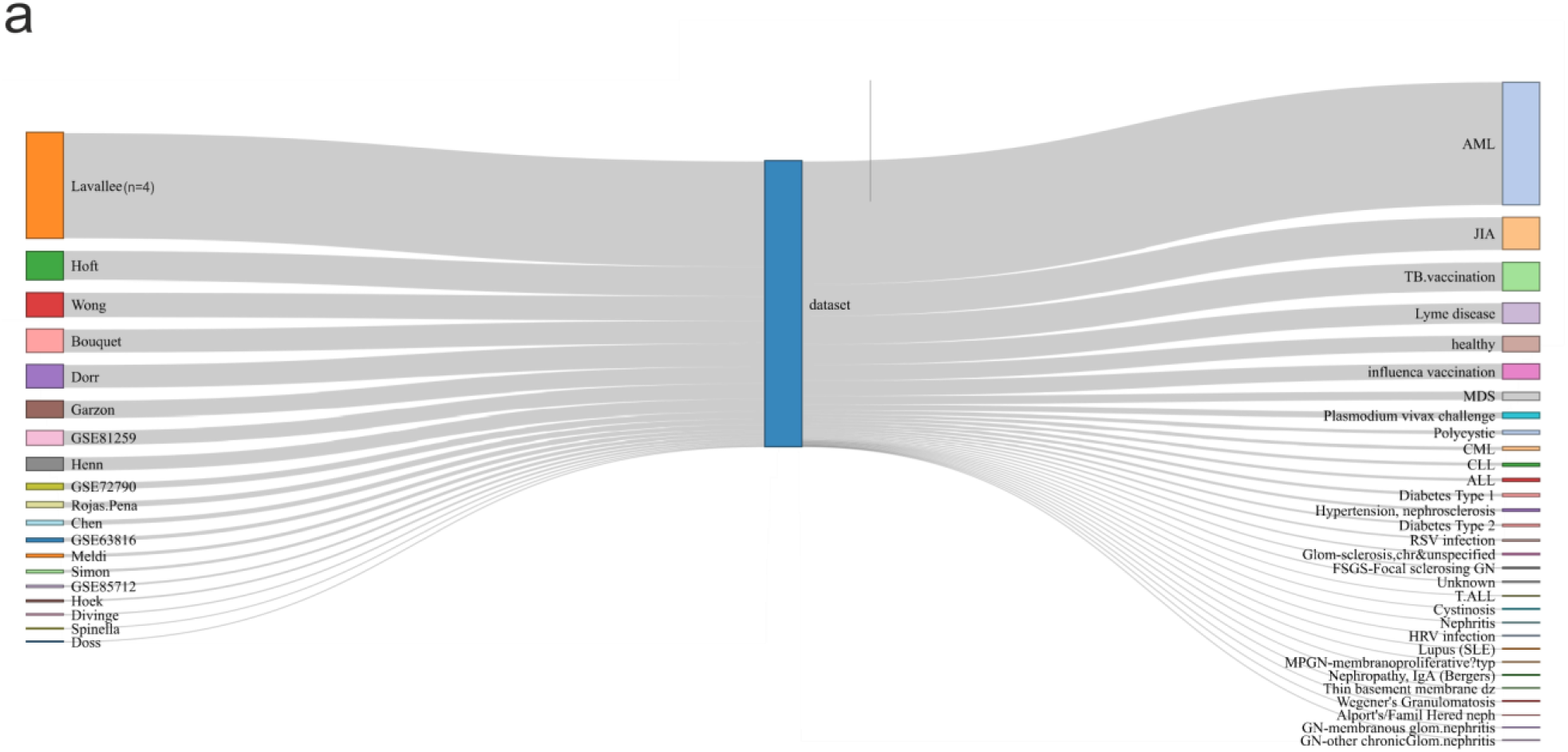
Overview of the studies used for evaluation. **a,** Sankey diagram of all 1.181 samples used in this study. Plot shows the link of each dataset to all represented diseases in this study. The width of the lines is proportionally to the flow quantity (individual studies (left), respectively diseases (right) within complete dataset).

**Extended Data Fig. 5:**
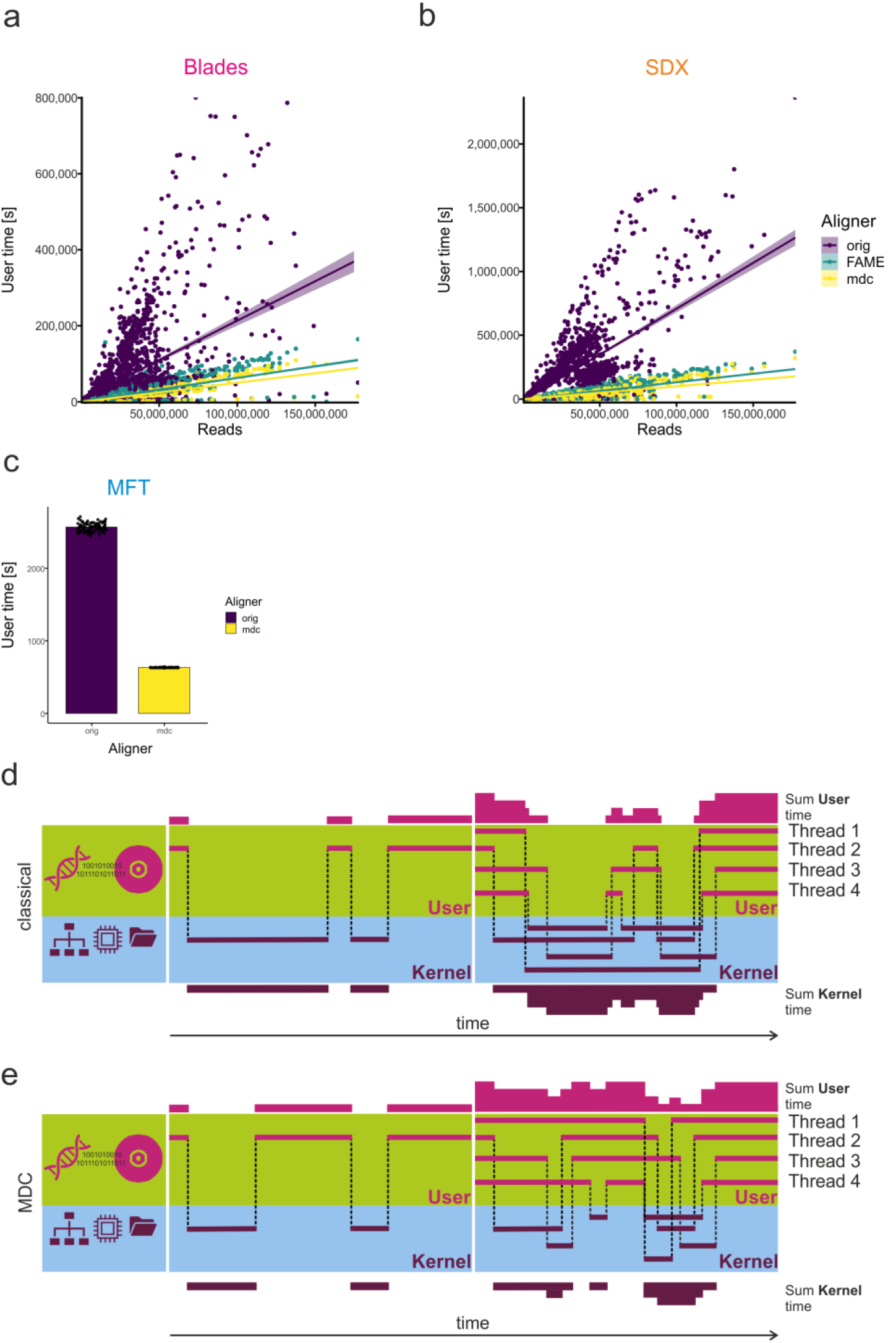
User time analysis of the pseudoalignment and quantification use case. **a, b,** Assessment of user times of 1.181 RNA-seq samples of different read sizes on Blades (**a**) and the SDX system (**b**). **c,** Comparison of user time for the ERR188140 data set. **d, e,** Depiction of user and kernel time for an application with one and three threads on a classical system (**d**), or with the use of MDC (**e**).

**Extended Data Fig. 6:**
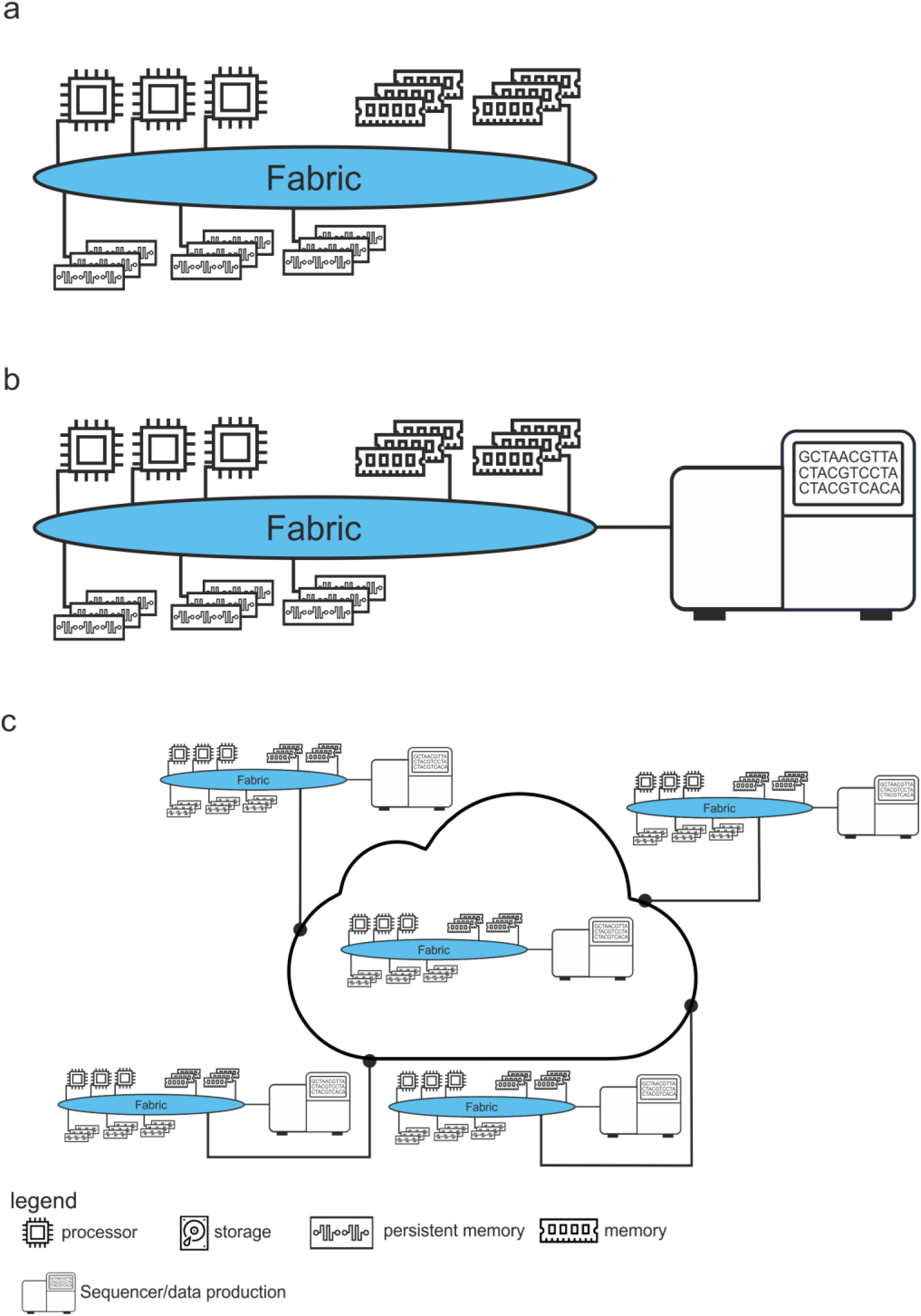
Scaling MDC to federated genomics processing. **a,** Architecture of MDC with central fabric. **b,** Sequencer directly attached to the MDC system. **c,** Federated cloud of MDC systems with connected sequencers.

**Extended Data Fig. 7:**
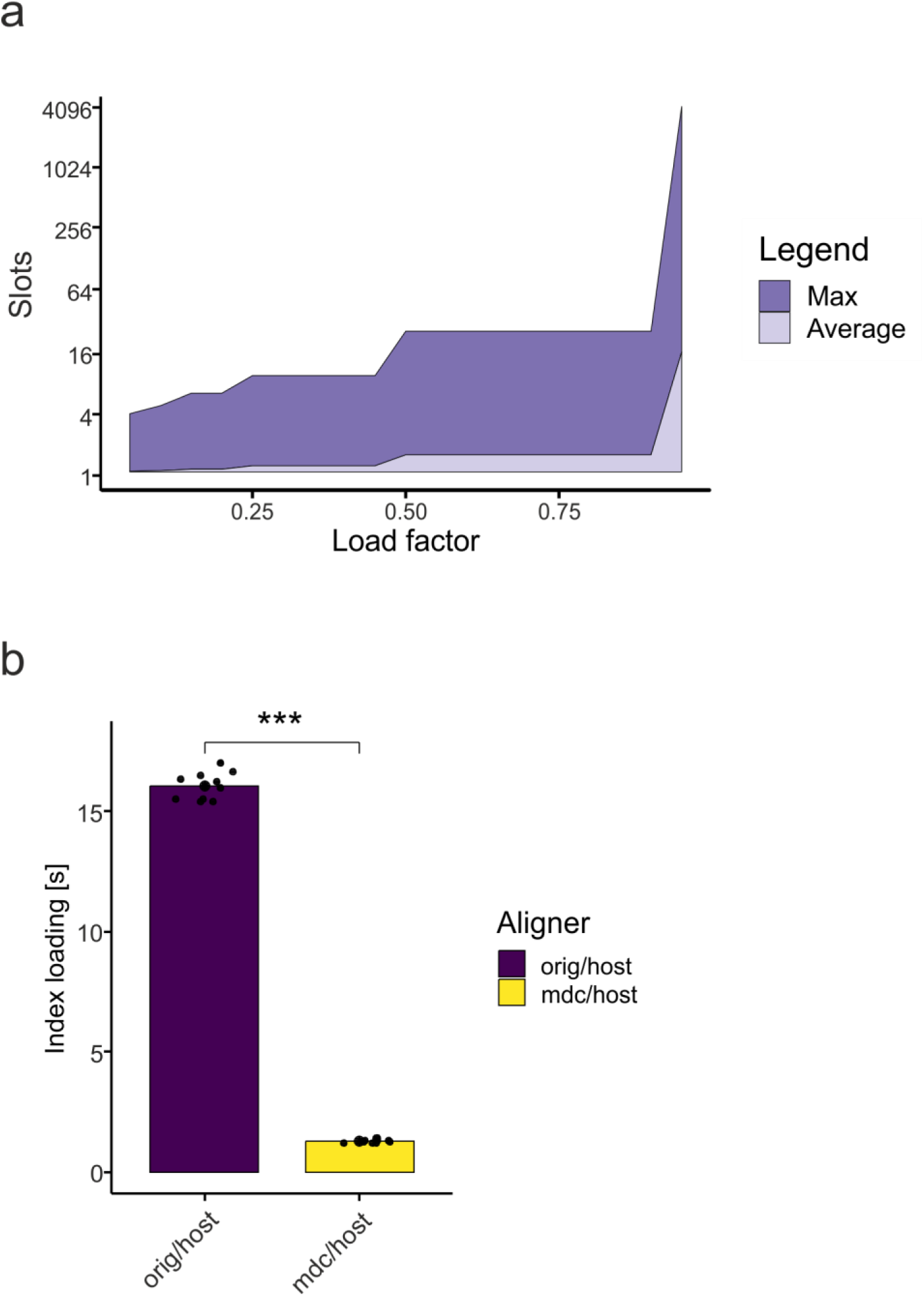
Kallisto version comparison. **a,** Average and maximum number of slots in the hash table that need to be searched per request with different load factors. **b,** Comparison of index loading times.

**Extended Data Fig. 8:**
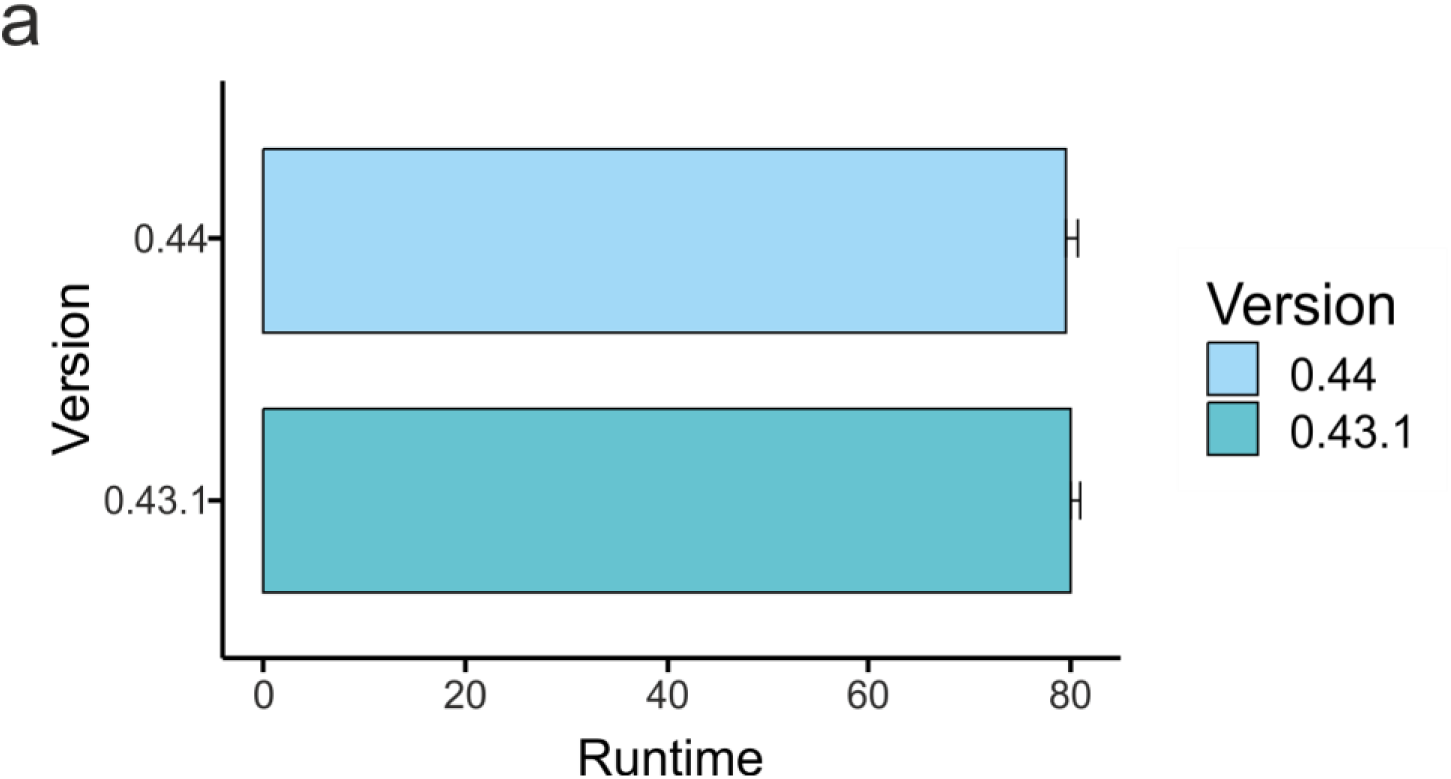
Hash table load factor impact. **a,** Runtime of the kallisto version used during preparation of the manuscript and the current version.

